# *lista*-GEM: the genome-scale metabolic reconstruction of *Lipomyces starkeyi*

**DOI:** 10.1101/2023.09.25.559328

**Authors:** Eduardo Almeida, Mauricio Ferreira, Wendel Silveira

## Abstract

Oleaginous yeasts cultivation in low-cost substrates is an alternative for more sustainable production of lipids and oleochemicals. *Lipomyces starkeyi* accumulates high amounts of lipids from different carbon sources, such as glycerol, and glucose and xylose (lignocellulosic sugars). Systems metabolic engineering approaches can further enhance its capabilities for lipid production, but no genome-scale metabolic networks have been reconstructed and curated for *L. starkeyi*. Herein, we propose *lista-*GEM, the first genome-scale metabolic model of *L. starkeyi*. We reconstructed the model using two high-quality models of oleaginous yeasts as templates and further curated the model to reflect the metabolism of *L. starkeyi*. We simulated phenotypes and predicted flux distributions in good accordance with experimental data. We also predicted targets to improve lipid production in glucose, xylose, and glycerol. The phase plane analysis indicated that the carbon availability affected lipid production more than oxygen availability. We found that the maximum lipid production in glucose and xylose required more oxygen than glycerol. Enzymes related to lipid synthesis in the endoplasmic reticulum were the main targets to improve lipid production: stearoyl-CoA desaturase, fatty-acyl-CoA synthase, diacylglycerol acyltransferase, and glycerol-3-phosphate acyltransferase. The glycolytic genes encoding pyruvate kinase, enolase, phosphoglycerate mutase, glyceraldehyde-3-phosphate dehydrogenase, and phosphoglycerate kinase were predicted as targets for overexpression. Pyruvate decarboxylase, acetaldehyde dehydrogenase, acetyl-CoA synthetase, adenylate kinase, inorganic diphosphatase, and triose-phosphate isomerase were predicted only when glycerol was the carbon source. Therefore, we demonstrated that *lista-*GEM provides multiple metabolic engineering targets to improve lipid production by *L. starkeyi* using carbon sources from agricultural and industrial wastes.

**Highlights:**

- *Lipomyces starkeyi* can accumulate high amounts of lipids from carbon sources found in agricultural and industrial wastes.
- We reconstructed *lista-*GEM, the first genome-scale metabolic model of *L. starkeyi*.
- Simulated phenotypes were in line with experimental results of *L. starkeyi*.
- We identified key gene targets for improving lipid production using metabolic engineering.

## Introduction

The optimization of bioprocesses allied with advances in genetic and metabolic engineering have enabled considerable advances in the production of lipids using oleaginous yeasts. These yeasts can accumulate at least 20% of their dry biomass as lipids, especially triacylglycerols (TAGs) (Salvador López et al., 2022). The requirement for sustainable sources of lipids for the production of oleochemicals, biodiesel, and human nutrition has boosted research with oleaginous yeasts able to use agricultural, industrial, and urban wastes as substrates for the development of bioprocesses (Abeln and Chuck, 2021; Spagnuolo et al., 2019).

*Lipomyces starkeyi* is an oleaginous yeast capable of growing and producing lipids using a diverse range of carbon sources, such as glucose, galactose, arabinose, xylose, glycerol, mannose, cellobiose, and sucrose (Smith and Kurtzman, 2011). Its growth and lipid production have been demonstrated in lignocellulosic biomasses, including corn stover (Pomraning et al., 2019), wheat straw (Yu et al., 2011), lignin derivatives (Putra et al., 2023); glycerol (Maruyama et al., 2018; Liu et al., 2017); and sewage sludge (Angerbauer et al., 2008). Importantly, *L. starkeyi* also tolerates inhibitors found in lignocellulosic hydrolysates, such as hydroxymethylfurfural, furfural, and phenolic compounds in synthetic media (Putra et al., 2023; Rahman et al., 2017) and detoxified wheat straw hydrolysate with high acetic acid concentration (4.2 g/L)(Yu et al., 2011). Besides, *L. starkeyi* can use levoglucosan, a major product from lignocellulose pyrolysis, as a carbon source (Ning et al., 2008).

Both lipid production and accumulation by *L. starkeyi* take place under nitrogen limitation conditions. This leads to an accumulation of citrate inside the cell, which is then converted to acetyl-CoA, kick-starting the fatty acid biosynthesis (Takaku et al., 2020). In contrast to other oleaginous yeasts, the malic enzyme of *L. starkeyi* can use both NAD^+^ and NADP^+^ as cofactors but prefers NAD^+^ (Tang et al., 2010). Thus, there is an additional requirement for NAD^+^ to drive fatty acid biosynthesis.

Efforts to engineer *L. starkeyi* strains can benefit from systems biology approaches related to genome-scale metabolic models (GEMs). These models aim to reconstruct and summarize the metabolic network of an organism based on its genome annotation, which is useful for both helping understand its physiology and for metabolic engineering endeavors (Ye et al., 2022). GEMs are mathematically formalized as an optimization problem, where a metabolic objective is maximized or minimized given the assumption of steady-state metabolism and constraints on the uptake of substrates and excretion of products (Orth et al., 2010). There have been many GEM reconstructions for oleaginous yeasts, such as *Yarrowia lipolytica* (Kavšček et al., 2015; Kerkhoven et al., 2016; Loira et al., 2012; Mishra et al., 2018; Pan and Hua, 2012; Wei et al., 2017), *Rhodotorula toruloides* (Dinh et al., 2019; Kim et al., 2021; Tiukova et al., 2019), *Papiliotrema laurentii* (Ventorim et al., 2022), and *Cutaneotrichosporon oleaginosus* (Pham et al., 2021). These models have been applied to predict essential genes and the use of different carbon and nitrogen sources, as well as better cultivation strategies and metabolic engineering targets. To the best of our knowledge, the GEM for *L. starkeyi* was still not reconstructed. Here, we present the first genome-scale reconstruction of *L. starkeyi*, termed *lista-*GEM. We show that *lista-*GEM successfully captures the growth and lipid-producing phenotype of *L. starkeyi* and, therefore, is a useful platform for *in silico* metabolic engineering of this yeast.

## Material and Methods

### Draft reconstruction and lipid metabolism

For the first draft of the genome-scale metabolic reconstruction of *L. starkeyi,* denominated *lista-*GEM, we used two well-curated GEMs as templates: *Y. lipolytica* iYali 4.1.2 (Kerkhoven et al., 2016) and *R*. *toruloides rhto*-GEM 1.3.0 (Tiukova et al., 2019). First, we identified the reactions from orthologs between the *L. starkeyi* genome NRRL Y-11557 (NCBI ID: 10576) and the *Y. lipolytica* or *R*. *toruloides* using bidirectional BLASTp (Madden, 2013). We considered as orthologs the genes with e-value < 1 x 10^-20^, identity > 35%, and alignment length > 150 bp. We excluded the reactions in iYali that were already present in *rhto*-GEM or that simplified lipid metabolism. Then, we retrieved the pseudo-reactions (e.g., biomass formation and exchange reactions) from *rhto*-GEM. We performed the reconstruction steps using the RAVEN Toolbox 2.7.9 (Wang et al., 2018) in MATLAB (The MathWorks Inc., Natick, Massachusetts).

To represent the lipid metabolism in *lista*-GEM, we used the Split Lipids Into Measurable Entities (SLIMEr) formalism (Sánchez et al., 2019), which describes lipids by splitting them into their basic components, such as pseudo-reactions that describe both the lipid classes and the acyl chain distributions. Here, we incorporated the following acyl chains of biotechnological importance: 16:0, 16:1, 18:0, 18:1, 18:2, 18.3.

### Biomass composition

From the total content of lipids, proteins, carbohydrates, RNA, and DNA retrieved from experimental measurements (Anschau et al., 2014; Matsuzawa et al., 2018; Probst and Vadlani, 2015), we updated the biomass equation of the *rhto*-GEM template and used it for *lista*-GEM. We also updated the biomass composition using data from glucose continuous cultures at a dilution rate of 0.06 h^-1^ (Anschau et al., 2014). We calculated the distribution of deoxyribonucleotides based on the GC content (47%) of *L. starkeyi* genome, as well as the sum of mRNAs and ncRNAs. For the amino acid distribution, we calculated it from the amino acid composition of translated coding sequences. We collected the contribution of triacylglycerols (TAGs), sterols, free FAs, phosphatidylcholine (PC), phosphatidylethanolamine (PE), phosphatidylinositol (PI), phosphatidylglycerol (PG), phosphatidylserine (PS), cardiolipin, and diacylglycerols (DAGs) from Probst and Vadlani (2015) and Uzuka et al. (1974). Considering data from Calvey et al. (2016), Matsuzawa et al. (2018), and Takaku et al. (2020), we adjusted the FA profile for the chains 16:0, 16:1, 18:0, 18:1, 18:2, and 18.3. The calculation procedures used to define the stoichiometric coefficients are provided in the *lista*-GEM documentation biomassCalculations.xlsx file available in the GitHub repository and Zenodo archive (See Data availability).

### Gap-filling, manual curation, and quality assessment

The gap-filling of *lista*-GEM was conducted in two steps. In the first step, we used Meneco (Prigent et al., 2017) to identify the reactions required for the biosynthesis of biomass components (target compounds) based on a list of available metabolites (seeds). The reactions identified by Meneco were retrieved from *rhto*-GEM. However, after Meneco was applied, we noticed that the model could still not sustain growth (i.e. produce biomass). Thus, in the second step, we used the “*fillGaps*” function from the RAVEN Toolbox. We considered growth on glucose (1 mmol/gDW h) at a biomass production rate of 0.01 h^-1^. The reactions required to sustain biomass formation were then retrieved from *rhto*-GEM and iYali templates. Finally, we noted that three reactions included from iYali (‘y300065’, ‘y300066’, ‘y200008’) were not required for growth and led to water and H^+^ overproduction in rich media simulations and removed them.

After the gap-filling step, we included the specific reactions required by *L. starkeyi* to sustain growth on the specified carbon sources and to meet cofactor requirements. In contrast to other oleaginous yeasts, the malic enzyme of *L. starkeyi* preferably uses NAD^+^ instead of NADP^+^ as a cofactor (Tang et al., 2010). Thus, we removed the malic enzyme reaction that used NADP^+^ and maintained only the one that uses NAD^+^. Additionally, we manually included the reactions necessary for L-rhamnose, lactose, cellobiose, and levoglucosan utilization. Finally, we updated the gene-reaction rules (grRules field), replacing the genes in the model that still contained the identification from the template (*R. toruloides*) with *L. starkeyi* homologs. The non-growth associated maintenance reaction remained the same as in *rhto*-GEM due to the lack of available data for *L. starkeyi*. We assessed the quality of the final reconstruction using MEMOTE (Lieven et al., 2020) (Figure 1A).

**Figure 1.**
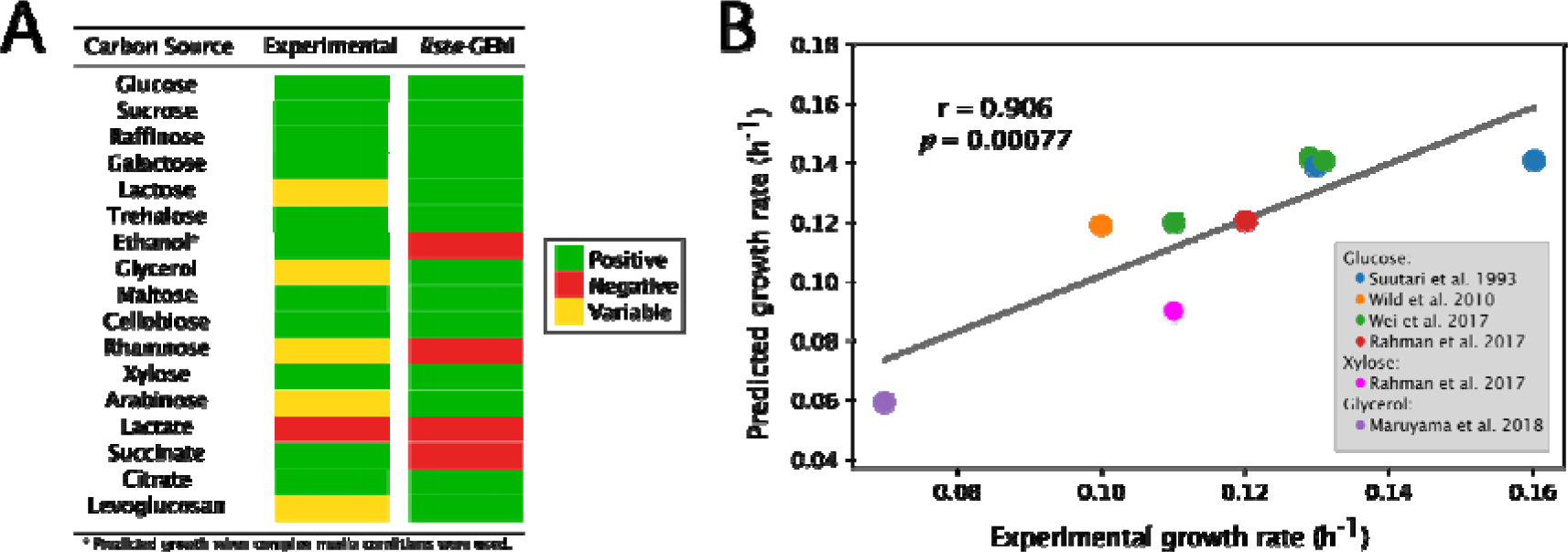
**(A)** Viability predicted by *lista*-GEM for different carbon sources compared to experimental data (Smith and Kurtzman, 2011). **(B)** Correlation between experimental and predicted growth rates on glucose, xylose and glycerol (Maruyama et al., 2018; Rahman et al., 2017; Suutari et al., 1993; Wild et al., 2010).

### Simulations and validation

To quantitatively assess the growth of *L. starkeyi* on different carbon sources (glucose, acetate, arabinose, cellobiose, citrate, ethanol, galactose, lactose, levoglucosan, xylose, rhamnose, R-lactate, S-lactate, mannose, trehalose; see Figure 1B) in minimal medium, we first constrained the lower bound of exchange reactions to zero and left only the oxygen, ammonium, H+, iron, phosphate, potassium, and sulfate exchange reactions unconstrained. Then, we allowed the uptake of each carbon source at -3 mmol/[g dry weight (DW) h] and optimized the formation of biomass via Flux Balance Analysis (FBA). To simulate growth on rich media, we also set the uptake of amino acids to -1 mmol/(g DW h).

To quantitatively assess model performance, we compared the experimental growth rate gathered from the literature with our predictions. Data were available for glucose, xylose and glycerol. When available, we set the carbon uptake rate as described in the manuscript. If not, we assumed a value of -3 mmol/(g DW h). The media (minimal or rich) was also adjusted based on the source manuscript description and the biomass formation was optimized using FBA. The correlation between *in vivo* and *in silico* growth data was determined by the Pearson’s correlation coefficient (Figure 3).

Furthermore, we conducted phase plane analysis in two different scenarios to determine conditions that would favor growth and lipid production in three carbon sources found in agro-industrial wastes (glucose and xylose from lignocellulosic biomasses and glycerol from biodiesel production). In the first scenario, we varied the carbon source uptake [from 0 to -10 mmol/(g DW h)] and oxygen uptake [from 0 to -50 mmol/(g DW h)] rates, while maintaining the other components of the minimal media as described above unconstrained. In the second scenario, instead of constraining carbon and oxygen uptake, we constrained nitrogen [from 0 to -9 mmol/(g DW h)] and oxygen [from 0 to -27 mmol/(g DW h)] uptake rates. We optimized the growth considering the biomass formation equation as described above, and simulated lipid production by optimizing a pseudoreaction representing TAG (1-16:0, 2-18:1, 3-18:1) exchange via FBA.

Moreover, we evaluated the main reactions related to lipid accumulation in nitrogen-limiting conditions using the environmental version of minimization of metabolic adjustment (eMOMA) (Kim et al., 2019). First, the same pseudoreaction described above to represent TAG exchange was added to the model, and the lower bound of the non-growth associated maintenance reaction (NGAM) was set to a low value [0.5 mmol/(g DW h)] to represent stationary growth. Then, we blocked the exchange reactions for ethanol, trehalose, butanediol, pyruvate, fumarate, 2-oxoglutarate, malate, oxaloacetate, glyoxylate, and acetate since we did not find evidence regarding the excretion of these metabolites for *L. starkeyi* under nitrogen-limiting conditions. We also blocked the exchange of decanoate, palmitate, palmitoleate, oleate, 14-demethyllanosterol, episterol, ergosterol, fecosterol, lanosterol, zymosterol, and ergosta-5,7,22,24(28)-tetraen-3beta-ol to promote TAG accumulation. Then, we set the growth as objective and performed FBA to obtain the flux distribution under non-restricted conditions (minimal media). Next, we blocked nitrogen exchange to simulate nitrogen restriction and confirmed that the model could not predict the growth and conducted the traditional MOMA between the model with and without nitrogen restriction. To test the reactions that affect lipid accumulation via knockout or overexpression, we removed reactions with zero flux in both conditions. Thereafter, we performed the eMOMA by knocking out or overexpressing (2x higher flux) the remaining reactions. We kept reactions where at least 2% increase in TAG exchange compared to the nitrogen-restricted reference and at least 90% growth remained compared to the nitrogen-abundant reference condition. We conducted eMOMA simulations for glucose, xylose, and glycerol at a fixed carbon uptake of -3 mmol/(g DW h).

Finally, we predicted overexpression targets to improve lipid production using glucose, xylose, and glycerol as carbon sources via flux scanning based on enforced objective flux (FSEOF) analysis(Choi et al., 2010). We performed the simulations considering minimal media, set the NGAM to 0 mmol/(g DW h), the TAG exchange pseudoreaction as the target, and the carbon uptake to -3 mmol/(g DW h). We conducted all simulations using the RAVEN Toolbox (v. 2.7.9) and/or the COBRA Toolbox (v. 3.4) (Heirendt et al., 2019) in MATLAB (The MathWorks Inc., Natick, Massachusetts) using Gurobi® (v. 10.0) as the solver.

## Results and Discussion

### Properties of the lista-GEM reconstruction

Herein, we reconstructed the first genome-scale metabolic model of the oleaginous yeast *Lipomyces starkeyi*. We applied a stepwise reconstruction strategy using the RAVEN toolbox based on the pipelines described by Tiukova et al. (2019) and Ventorim et al. (2022). Most genes (907 of 935) in the model were recovered in the first step (Homology draft; Table 1) of the reconstruction via bidirectional BLAST with the *R. toruloides* and *Y. lipolytica*, and their respective GEMs (*rhto*-GEM 1.3.0 and iYali 4.1.2). The next steps of the reconstruction focused mainly on adding pseudo and lipid metabolism (SLIMEr) reactions, gap-filling and manual curation of the model (see the Material and Methods). The final version of the model *lista*-GEM 1.0.0 presented a final MEMOTE score of 52%. This low stoichiometry consistency is related to the fact that the model was penalized for stoichiometric consistency and annotation, a common phenomenon for models that included the SLIMEr formalism, as lipid species are normalized by their weight for direct integration of lipid measurements (Sánchez et al., 2019). Consistently, the GEMs *rhto-*GEM (Tiukova et al., 2019) and *papla*-GEM (Ventorim et al., 2022), which included the SLIMEr formalism, have a score similar to *lista*-GEM.

**Table 1.**
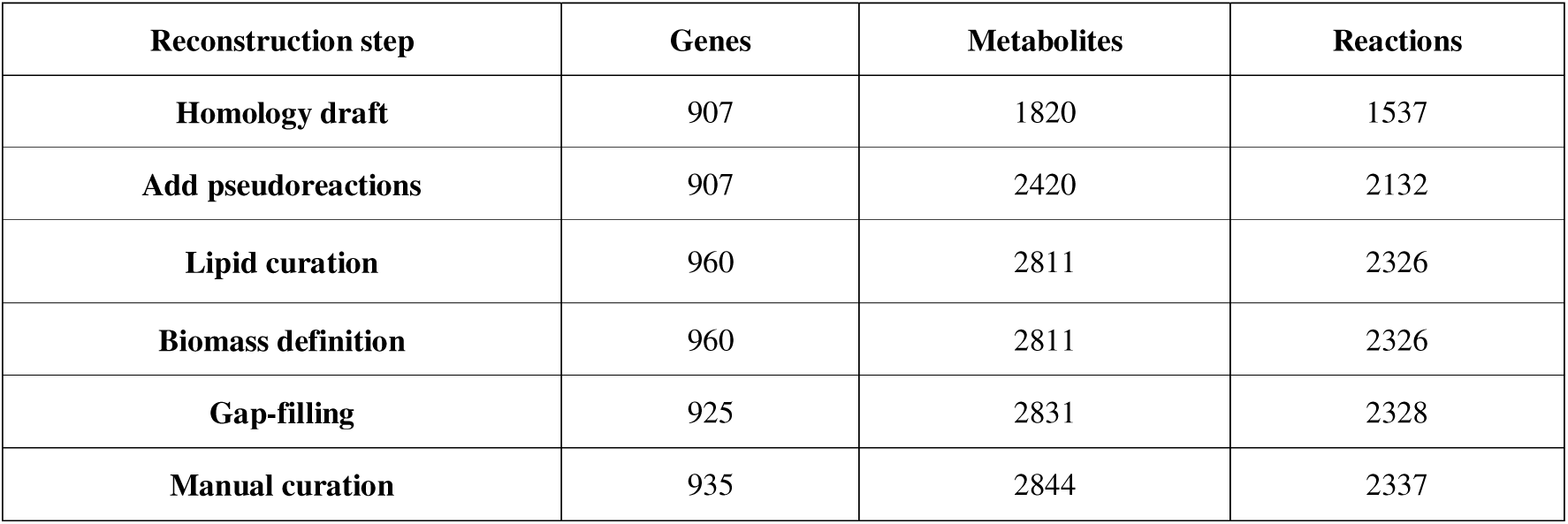
Genes, metabolites, and reactions of *lista*-GEM during the reconstruction steps.

| Reconstruction step | Genes | Metabolites | Reactions |
| --- | --- | --- | --- |
| Homology draft | 907 | 1820 | 1537 |
| Add pseudoreactions | 907 | 2420 | 2132 |
| Lipid curation | 960 | 2811 | 2326 |
| Biomass definition | 960 | 2811 | 2326 |
| Gap-filling | 925 | 2831 | 2328 |
| Manual curation | 935 | 2844 | 2337 |

### lista-GEM accurately represents the metabolism of L. starkeyi

The model presented a good performance for qualitative and quantitative growth prediction compared to experimental data. For 14 of 17 carbon sources, the model correctly predicted the growth/non-growth profile (Figure 1A) (Smith and Kurtzman, 2011). For ethanol, growth was predicted only in complex media simulations with uptake of essential amino acids. However, the model could not predict growth in none of the conditions tested for succinate. Furthermore, the predicted growth rates in three carbon sources of biotechnological interest (glucose, xylose, and glycerol) presented a good Pearson correlation with *in vivo* measurements (r = 0.906, *p =* 0.00077) (Maruyama et al., 2018; Rahman et al., 2017; Suutari et al., 1993; Wild et al., 2010) (Figure 1B).

The phase plane analysis for growth and TAG production in glucose, xylose and glycerol as carbon sources indicated a higher dependence on carbon uptake than oxygen uptake (Figures 2-3, S1-4). Although, a minimum oxygen uptake [15-20 mmol/(g DW h) for glucose and xylose and 5-10 mmol/(g DW h) for glycerol] was required by *L. starkeyi* to reach maximum biomass production, further increases in oxygen availability, in contrast to carbon availability, did not increased biomass production.

**Figure 2.**
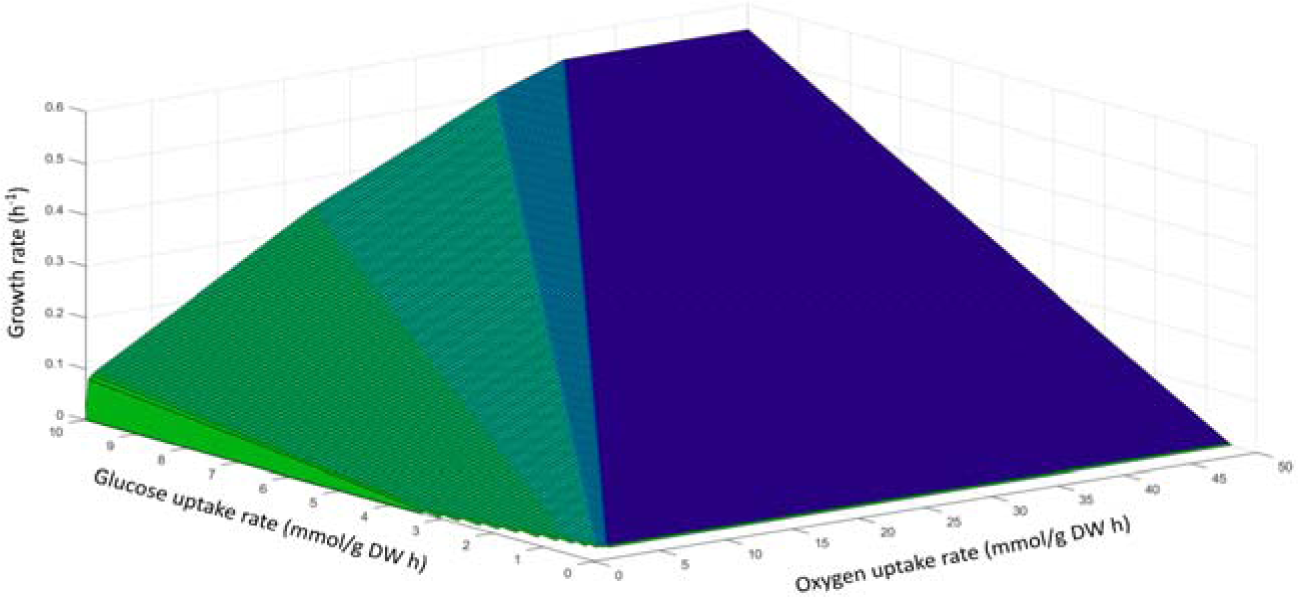
Effects of varying the glucose and oxygen uptake rates on growth.

**Figure 3.**
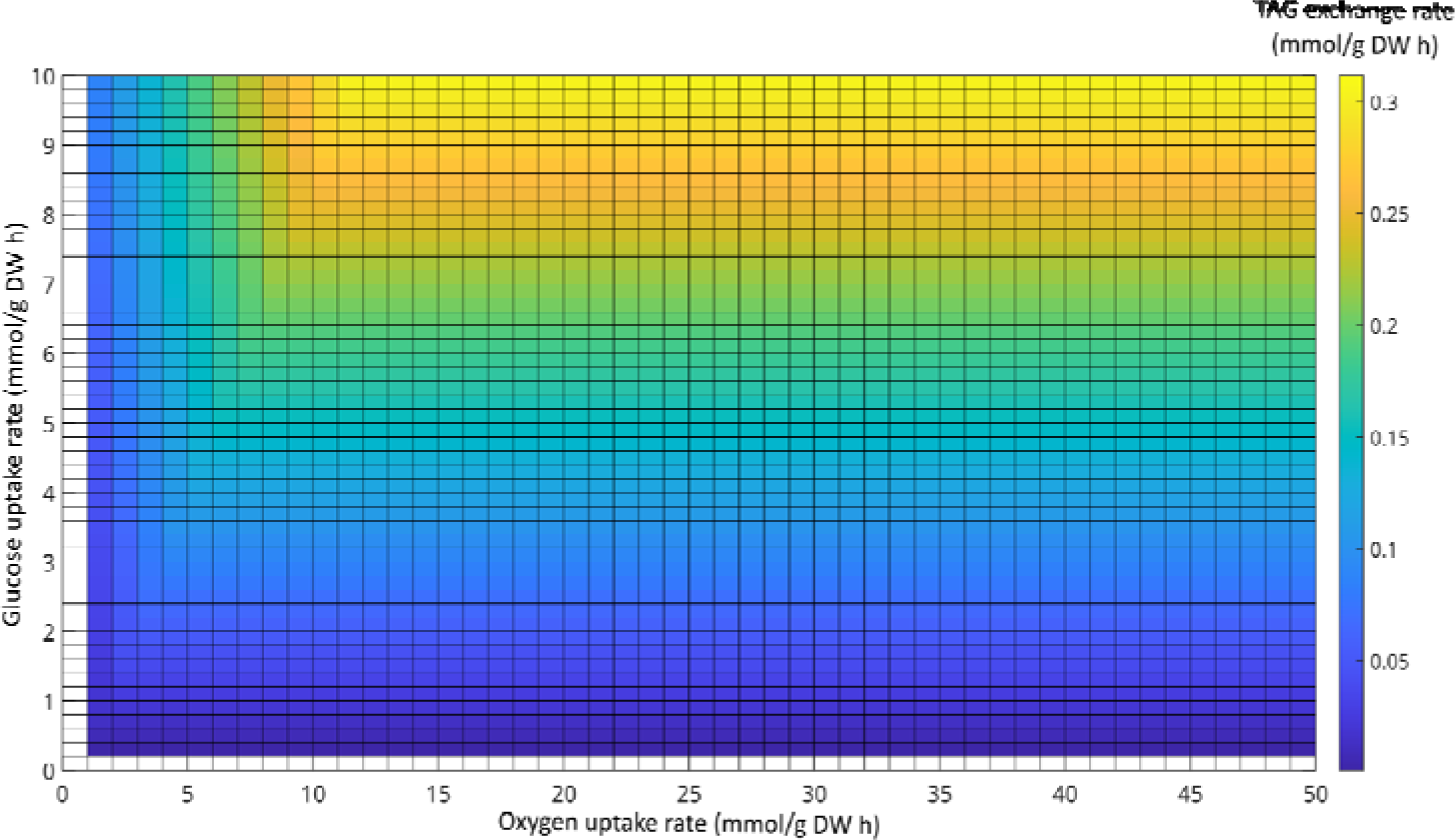
Effects of varying the glucose and oxygen uptake rates on TAG exchange.

Notably, the oxygen requirement for maximum TAG production was lower than those for growth on glucose and xylose (Figures 1-2 and S1-2). This is likely associated with the fact that lipid accumulation in oleaginous yeasts starts from the late-exponential phase (Ratledge, 2008), where the oxygen availability is lower than the exponential phase. Importantly, from a bioprocess development point of view, the low oxygen requirement for TAG accumulation in *L. starkeyi* is advantageous, as the dissolved oxygen availability, which in turn affects the oxygen transfer rate, is a limiting factor for aerobic processes. Otherwise, the oxygen requirement for maximum growth and TAG production was the same on glycerol (Figures S3-4), highlighting important differences between glycolytic and gluconeogenic carbon sources regarding the lipid production by *L. starkeyi*. For the phase plane analysis performed varying the nitrogen source uptake rate and oxygen uptake rate, we noticed that the key determinant for achieving growth and producing TAG was the availability of oxygen (Figures S5-7).

### Predicted targets for enhancing the production of TAGs by metabolic engineering strategies

To improve the production of lipids using metabolic engineering, we identified gene targets for knockout or overexpression using a combination of minimal adjustment of fluxes (eMOMA) and flux scanning (FSEOF). We simulated growth conditions where nitrogen was limited and the NGAM was set to a low value, which represents stationary growth, and used a TAG representative to simulate lipid production (see Material and Methods).

The eMOMA approach is useful to predict the flux distribution for a changed environment. Similar to MOMA, eMOMA is implemented as a linear or quadratic problem to minimize the L1 or L2-norm distances, respectively, between the reference and alternative flux distributions. However, while the MOMA approach is tailored to minimize the difference between a wild-type and a mutant strain based on the principle of minimal metabolic adjustment, the eMOMA approach expands the MOMA implementation by considering an additional constraint, where the flux through the uptake reaction of a growth-limiting nutrient is equal to zero. To predict the important reactions for lipid production, we constrained the uptake of exchange reactions relative to metabolites from the TCA cycle, sterols, and various lipids such as decanoate, palmitate, palmitoleate and oleate. This approach allowed the identification of the main reactions associated with lipid accumulation. It is important to point out that many of the identified reactions were the same in glucose, xylose, or glycerol as the carbon sources, being primarily involved in the exchange reaction of the carbon source, the pentose phosphate pathway, tricarboxylic acid (TCA) cycle, and lipid metabolism (Tables 2-4).

**Table 2.**
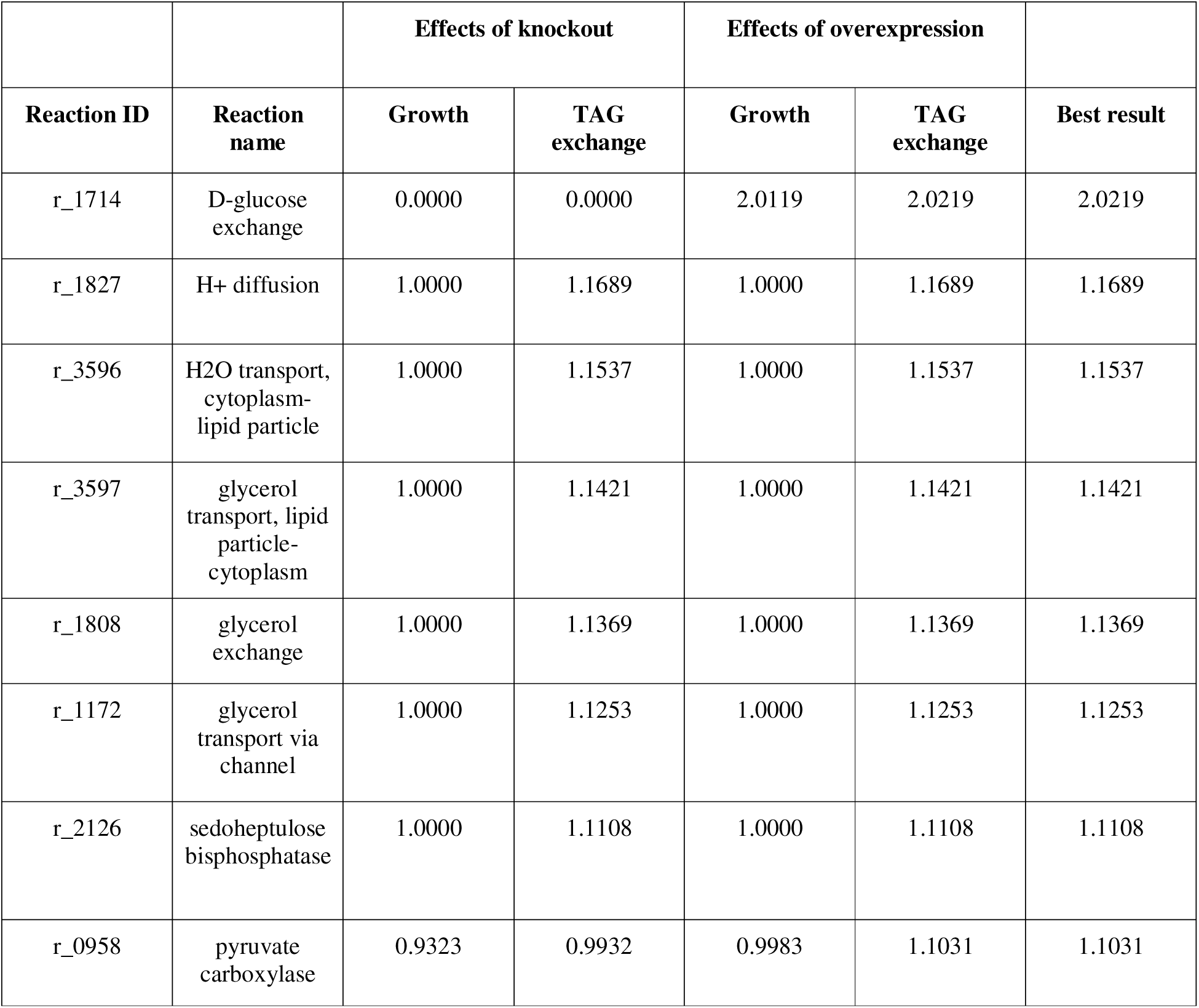
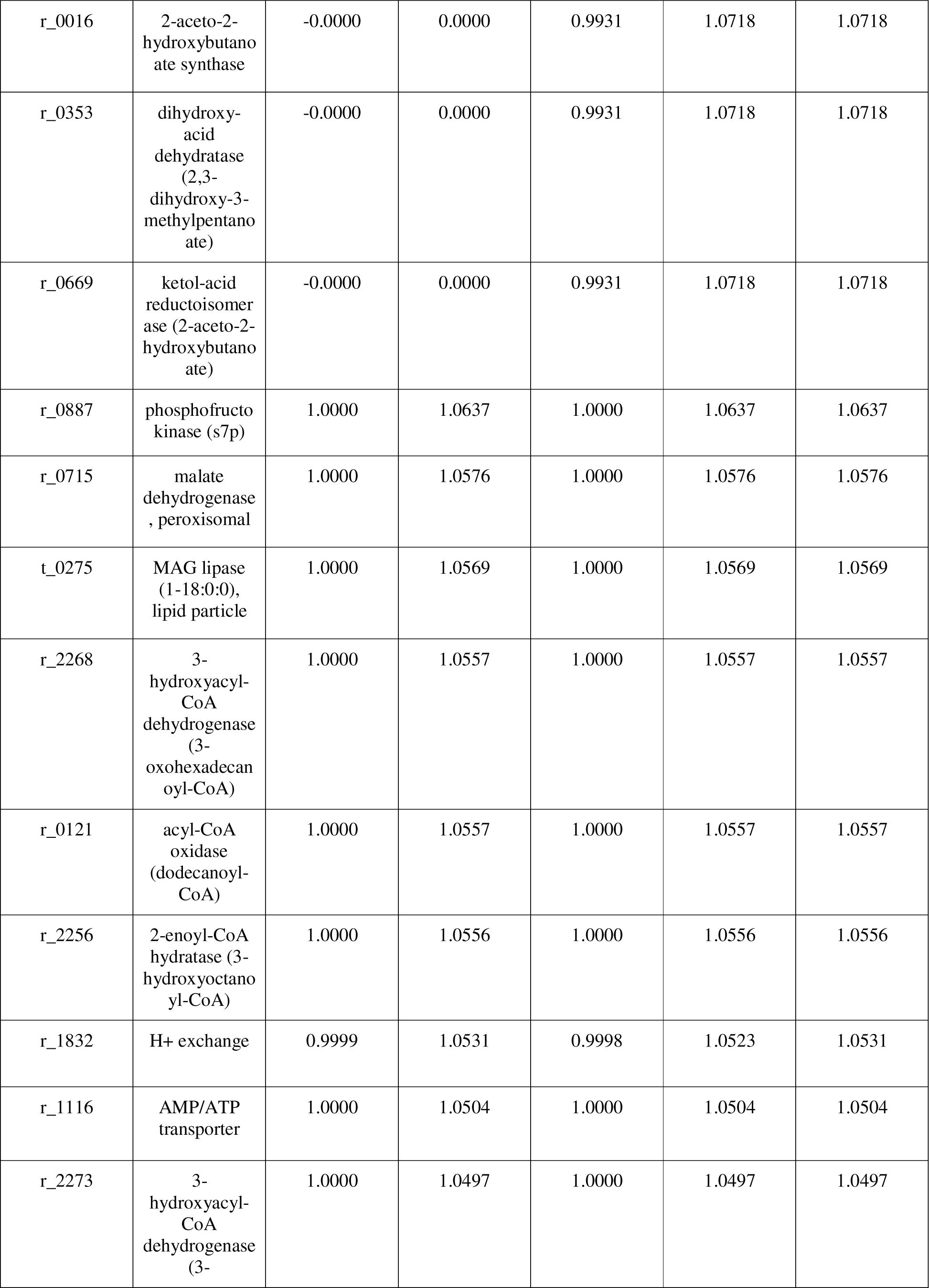
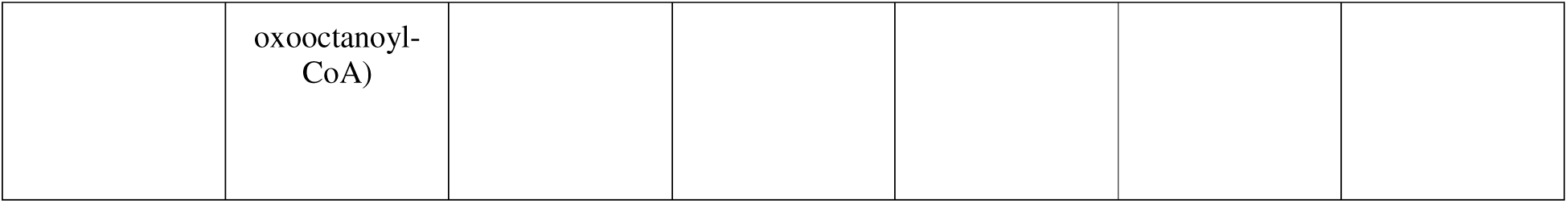
Top 20 targets identified by eMOMA on *lista-*GEM using glucose as carbon source.

**Table 3.** Top 20 targets identified by eMOMA on *lista-*GEM using xylose as carbon source.

|  |  | Effects of knockout |  | Effects of overexpression |  |  |
| --- | --- | --- | --- | --- | --- | --- |
| Reaction ID | Reaction name | Growth | TAG exchange | Growth | TAG exchange | Best result |
| r_1718 | D-xylose exchange | 0.0000 | 0.0000 | 2.0145 | 2.0308 | 2.0308 |
| r_2126 | sedoheptulose biphosphatase | 1.0000 | 1.1539 | 1.0000 | 1.1539 | 1.1539 |
| r_0887 | phosphofructokinase (s7p) | 1.0000 | 1.0869 | 1.0000 | 1.0869 | 1.0869 |
| r_0016 | 2-aceto-2-hydroxybutanoate synthase | -0.0000 | 0.0055 | 0.9931 | 1.0804 | 1.0804 |
| r_0353 | dihydroxy-acid dehydratase (2,3-dihydroxy-3-methylpentanoate) | -0.0000 | 0.0055 | 0.9931 | 1.0804 | 1.0804 |
| r_0669 | ketol-acid reductoisomerase (2-aceto-2-hydroxybutanoate) | -0.0000 | 0.0055 | 0.9931 | 1.0804 | 1.0804 |
| r_0958 | pyruvate carboxylase | 0.9323 | 0.9505 | 0.9983 | 1.0795 | 1.0795 |
| r_1980 | O2 transport | 1.0000 | 1.0637 | 1.0000 | 1.0637 | 1.0637 |
| r_0715 | malate dehydrogenase, peroxisomal | 1.0000 | 1.0562 | 1.0000 | 1.0562 | 1.0562 |
| r_1930 | malate/oxaloacetate shuttle | 1.0000 | 1.0554 | 1.0000 | 1.0554 | 1.0554 |
| r_0468 | glutamate 5-kinase | 1.0000 | 1.0539 | 1.0000 | 1.0539 | 1.0539 |
| r_0473 | glutamate-5-semialdehyde dehydrogenase | 1.0000 | 1.0539 | 1.0000 | 1.0539 | 1.0539 |
| r_1116 | AMP/ATP transporter | 1.0000 | 1.0490 | 1.0000 | 1.0490 | 1.0490 |
| r_2256 | 2-enoyl-CoA hydratase (3-hydroxyoctanoyl-CoA) | 1.0000 | 1.0488 | 1.0000 | 1.0488 | 1.0488 |
| r_2273 | 3-hydroxyacyl-CoA dehydrogenase (3-oxooctanoyl-CoA) | 1.0000 | 1.0488 | 1.0000 | 1.0488 | 1.0488 |
| r_2285 | acetyl-CoA C-acyltransferase (hexanoyl-CoA) | 1.0000 | 1.0488 | 1.0000 | 1.0488 | 1.0488 |
| r_1832 | H<sup>+</sup> exchange | 0.9999 | 1.0333 | 0.9998 | 1.0444 | 1.0444 |
| r_2268 | 3-hydroxyacyl-CoA dehydrogenase (3-oxohexadecanoyl-CoA) | 1.0000 | 1.0443 | 1.0000 | 1.0443 | 1.0443 |
| r_2251 | 2-enoyl-CoA hydratase (3-hydroxyhexadecanoyl-CoA) | 1.0000 | 1.0443 | 1.0000 | 1.0443 | 1.0443 |
| r_0107 | acetyl-CoA C-acyltransferase (decanoyl-CoA) | 1.0000 | 1.0438 | 1.0000 | 1.0438 | 1.0438 |

**Table 4.** Top 20 targets identified by eMOMA on *lista-*GEM using glycerol as carbon source.

| Reaction ID | Reaction name | Effects of knockout |  | Effects of overexpression |  | Best result |
| --- | --- | --- | --- | --- | --- | --- |
|  |  | Growth | TAG exchange | Growth | TAG exchange |  |
| r_1808 | glycerol exchange | 0.0000 | 0.0000 | 2.0249 | 2.0245 | 2.0245 |
| r_0958 | pyruvate carboxylase | 0.9384 | 1.0884 | 0.9980 | 1.1180 | 1.1180 |
| r_0353 | dihydroxy-acid dehydratase (2,3-dihydroxy-3-methylpentanoate) | -0.0000 | 0.0000 | 0.9930 | 1.1058 | 1.1058 |
| r_0669 | ketol-acid reductoisomerase (2-aceto-2-hydroxybutanoate) | -0.0000 | 0.0000 | 0.9930 | 1.1058 | 1.1058 |
| r_0016 | 2-aceto-2-hydroxybutanoate synthase | -0.0000 | 0.0000 | 0.9930 | 1.1058 | 1.1058 |
| r_0219 | aspartate-semialdehyde dehydrogenase | -0.0000 | 0.0000 | 0.9919 | 1.0668 | 1.0668 |
| r_0215 | aspartate kinase | -0.0000 | 0.0000 | 0.9919 | 1.0668 | 1.0668 |
| r_1832 | H<sup>+</sup> exchange | 0.9861 | 1.0198 | 1.0000 | 1.0604 | 1.0604 |
| r_1794 | formate transport | 0.9999 | 1.0476 | 1.0000 | 1.0471 | 1.0476 |
| r_2045 | serine transport | 0.9999 | 1.0476 | 1.0000 | 1.0472 | 1.0476 |
| r_0503 | glycine hydroxymethyltransferase | 0.9999 | 1.0476 | 1.0000 | 1.0472 | 1.0476 |
| r_0724 | methenyltetrahydrikate cyclohydrolase | 0.9999 | 1.0476 | 1.0000 | 1.0471 | 1.0476 |
| r_0733 | methylenetetrahydrofolate dehydrogenase (NADP) | 0.9999 | 1.0476 | 1.0000 | 1.0430 | 1.0476 |
| r_0447 | formate-tetrahydrofolate ligase | 0.9999 | 1.0476 | 1.0000 | 1.0471 | 1.0476 |
| r_1811 | glycine transport | 0.9999 | 1.0473 | 1.0000 | 1.0463 | 1.0473 |
| r_2141 | fatty-acyl-CoA synthase (n-C18:0CoA) | 1.0000 | 0.9860 | 0.9913 | 1.0444 | 1.0444 |
| r_0725 | methenyltetrahydrofolate cyclohydrolase | 0.9999 | 1.0438 | 0.9977 | 0.0000 | 1.0438 |
| r_0732 | methylenetetrahydrofolate dehydrogenase (NADP) | 1.0000 | 1.0421 | 0.2523 | 0.2539 | 1.0421 |
| r_0446 | formate-tetrahydrofolate ligase | 1.0000 | 1.0393 | 1.0000 | 1.0393 | 1.0393 |

The FSEOF approach relies on enforcing the flux on product formation by searching for candidate reactions whose increase in flux also increases the flux on product formation. This ensures that all identified reactions contribute to enhancing the formation of the desired product. The maximum theoretical value for product formation predicted by conventional FBA is biologically unrealistic since the formation of biomass becomes negligible. To circumvent this, FSEOF sets as the objective function the biomass formation and identifies the intracellular fluxes that increase when the maximum theoretical value for product formation is applied as a constraint. This makes it possible to achieve a product formation flux close to the maximum theoretical value while respecting biological feasibility. The FSEOF analysis identified similar reactions to eMOMA, such as those involved in pyruvate metabolism, the pentose phosphate pathway, and TCA cycle. Additionally, we also identified reactions such as acetyl-CoA carboxylase, acetyl-CoA synthetase, and fatty-acyl-CoA synthase, which are key reactions involved in the biosynthesis of fatty acids (Table 5).

**Table 5.** Targets identified by FSEOF to improve lipid production using *lista-*GEM.

| % of remaining TAG production |  |  |  |  |
| --- | --- | --- | --- | --- |
| Glucose | Xylose | Glycerol | Reaction name | Genes |
| 18.21 | 18.17 | 19.26 | pyruvate kinase | ODQ76269.1 |
| 17.99 | 17.96 | 18.67 | acetyl-CoA carboxylase | ODQ75673.1 and ODQ72018.1 |
| 17.76 | 17.73 | 18.88 | enolase | ODQ74427.1 |
| 17.76 | 17.73 | 18.88 | phosphoglycerate mutase | ODQ73127.1 or ODQ76545.1 |
| 17.28 | 17.24 | 18.46 | glyceraldehyde-3-phosphate dehydrogenase | ODQ75822.1 |
| 17.28 | 17.24 | 18.46 | phosphoglycerate kinase | ODQ69690.1 |
| 16.75 | 16.72 | 17.60 | bicarbonate formation | ODQ71959.1 |
|  |  | 21.09 | pyruvate decarboxylase | ODQ73413.1 |
|  |  | 21.09 | acetaldehyde dehydrogenase | ODQ69191.1 or ODQ69510.1 or ODQ70419.1 or ODQ70788.1 or ODQ71777.1 or ODQ72007.1 or ODQ73381.1 or ODQ75322.1 |
|  |  | 20.98 | acetyl-CoA synthetase | ODQ74542.1 |
|  |  | 18.90 | adenylate kinase | ODQ72719.1 or ODQ74535.1 |
|  |  | 17.24 | inorganic diphosphatase | ODQ71884.1 |
|  |  | 16.45 | triose-phosphate isomerase | ODQ69158.1 or ODQ69323.1 |
| 1.67 | 1.67 | 1.72 | stearoyl-CoA desaturase (n-C18:0CoA -> n-C18:1CoA), ER membrane | ODQ73471.1 |
| 1.65 | 1.65 | 1.69 | fatty-acyl-CoA synthase (n-C18:0CoA) | ODQ70131.1 or (ODQ75304.1 and ODQ70130.1) or ODQ70131.1 or ODQ75303.1 or ODQ75304.1 |
| 0.89 | 0.89 | 0.92 | diacylglycerol acyltransferase (1-16:0, 2-18:1, 3-18:1), endoplasmic reticulum membrane | ODQ70106.1 or ODQ75508.1 |
| 0.82 | 0.82 | 0.85 | PA phosphatase (1-16:0, 2-18:1),<br>endoplasmic reticulum membrane | ODQ71343.1 |
| 0.82 | 0.82 | 0.85 | 1-acyl-sn-glycerol-3-phosphate<br>acyltransferase (1-16:0, 2-18:1),<br>endoplasmic reticulum membrane | ODQ69760.1 or ODQ75695.1 |
| 0.79 | 0.79 | 0.82 | glycerol-3-phosphate acyltransferase<br>(16:0), endoplasmic reticulum<br>membrane | ODQ74991.1 |
| 0.69 | 0.69 | 0.73 | fatty-acyl-CoA synthase (n-C16:0CoA) | ODQ70131.1 or (ODQ75304.1 and<br>ODQ70130.1) or ODQ70131.1 or<br>ODQ75303.1 or ODQ75304.1 |
|  |  | 0.79 | glycerol kinase | ODQ76330.1 |
|  | 0.76 |  | glycerol-3-phosphate dehydrogenase<br>(NAD) | ODQ74980.1 |

Similar to other metabolic engineering strategies reported for other oleaginous yeasts, we could identify common targets for overexpression and knockout. The eMOMA and FSEOF analyses performed on the *rhto-*GEM and *i*MK735 (Kim et al., 2019) models also predicted many of the same targets shown on Tables 2-5. Additionally, many of the target genes predicted using *lista-*GEM are experimentally validated for *L. starkeyi* using the three tested sugars, such as the acetyl-CoA carboxylase, fatty acid synthetase and glycerol-3-phosphate acyltransferase (Zhang et al., 2022). The overexpression of these genes has been performed by Jeffries et al. 2017 (Jeffries et al., 2017) to improve lipid production using stillage and its derivatives. They reported an increase of 85% in the lipid titer. In another study, the overexpression of the gene coding for acetyl-CoA carboxylase led to an increase in the production of malonyl-CoA (Lu et al., 2008). Further, by doubling the copy number of the genes encoding subunits of the enzyme fatty-acid synthetase, Chen et al. (2020) reported an increase of 60% in the lipid content. Taken together, our results highlight the accuracy of the *lista-*GEM reconstruction to identify important reactions for lipid production and to develop suitable metabolic engineering strategies to enhance lipid production.

## Conclusions

Herein, we present the first genome-scale metabolic reconstruction and curation of *L. starkeyi*, termed *lista*-GEM. The model was based on two high-quality reconstructions and further curated using experimental data to represent the metabolic specificities of *L. starkeyi*. The growth conditions simulated using FBA were in good agreement with experimental growth data, underscoring the usefulness of *lista*-GEM in predicting phenotypes. Further, its usefulness for metabolic engineering was demonstrated by the prediction of gene targets in line with experimental results. Although a genome-scale reconstruction only describes current knowledge and is never finished (Anton et al., 2023), the open nature of GEMs allows for its continuous development by all members of the scientific community. While the *lista*-GEM already proves itself useful for the study of lipid metabolism and for biotechnological applications, enhancements such as enzyme constraints can further push it to new horizons.

## Data availability

The scripts used for the reconstruction and simulations as well as the *lista*-GEM model are available in the GitHub repository at https://github.com/LabFisUFV/lista-GEM or through Zenodo at https://doi.org/10.5281/zenodo.8367982. The model is provided according to the standard-GEM template (Anton et al., 2023) and in different formats (TXT, SMBL, XLSX, and MAT).

## Funding statement

This study was financed in part by the Coordenação de Aperfeiçoamento de Pessoal de Nível Superior – Brasil (CAPES) [Finance Code 001]. MF acknowledges funding from CAPES. EA acknowledges funding from Conselho Nacional de Desenvolvimento Científico e Tecnológico – Brasil (CNPq) [Finance Code 140538/2021-6]. WS acknowledges funding from CNPq [Finance Code 312390/2020-3]. The funders had no role in study design, data collection and analysis, decision to publish, or preparation of the manuscript.

## Conflicts of interest

The authors declare no competing interests.

## Supplementary material

**Figure S1.**
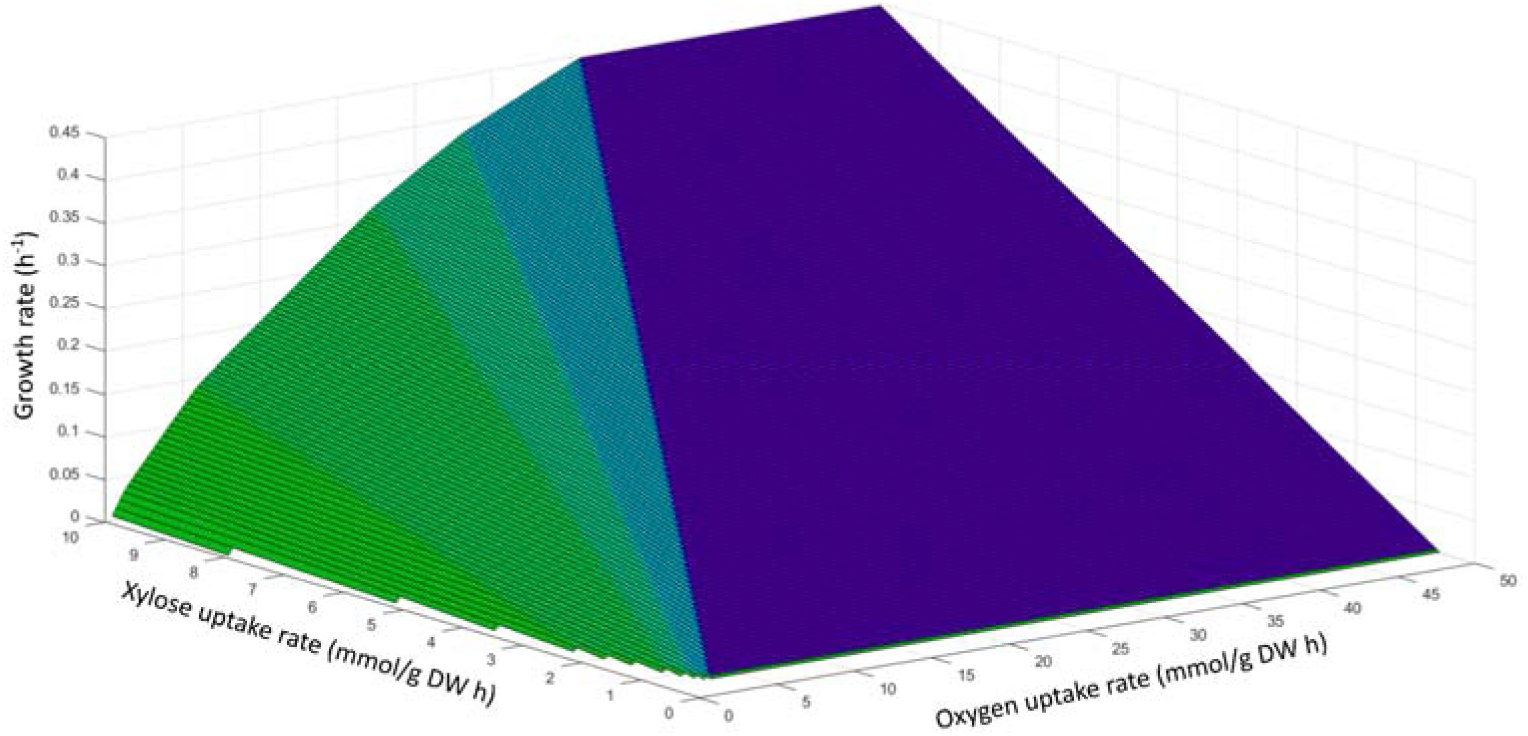
Effects of varying the xylose and oxygen uptake rates on growth.

**Figure S2.**
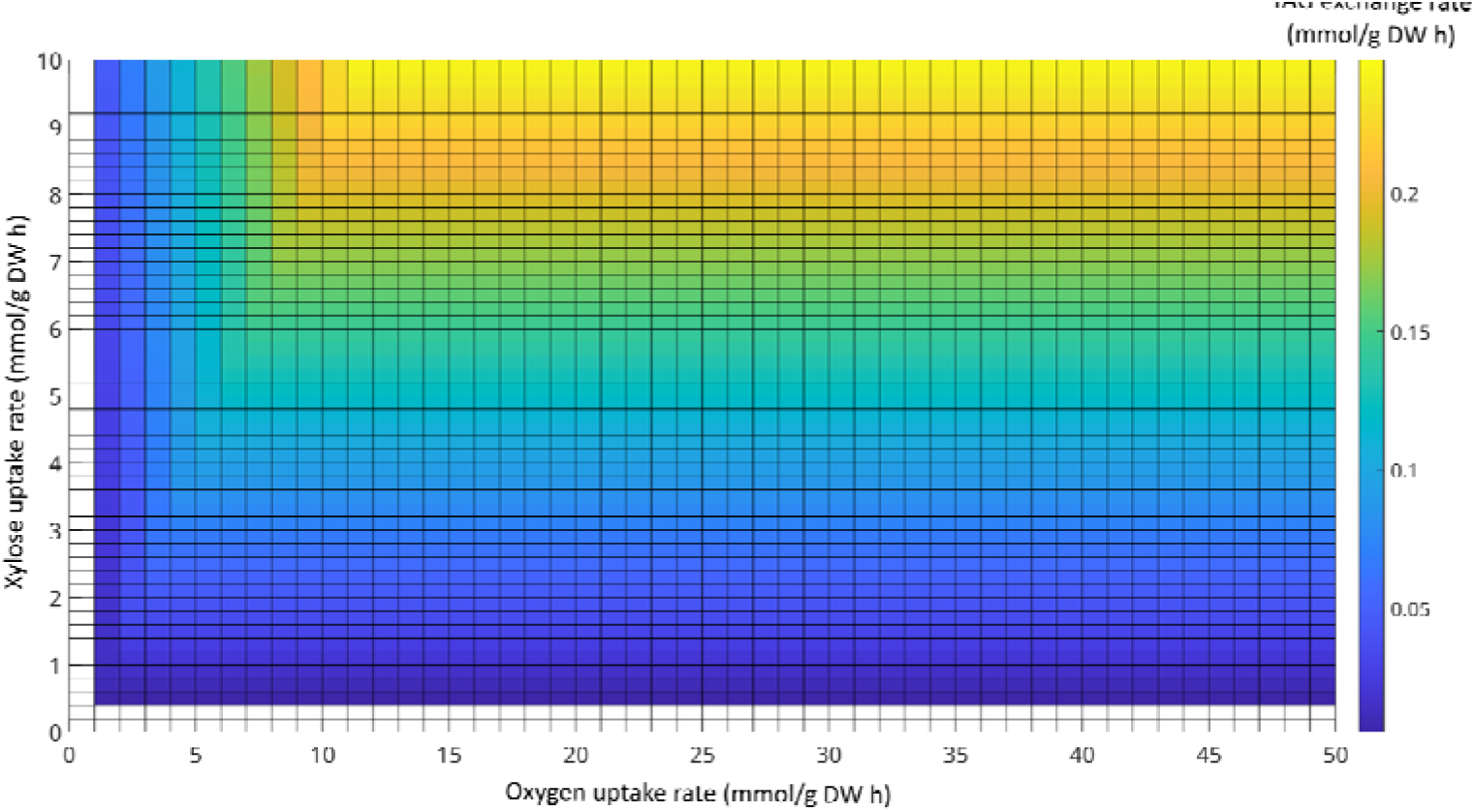
Effects of varying the xylose and oxygen uptake rates on TAG exchange.

**Figure S3.**
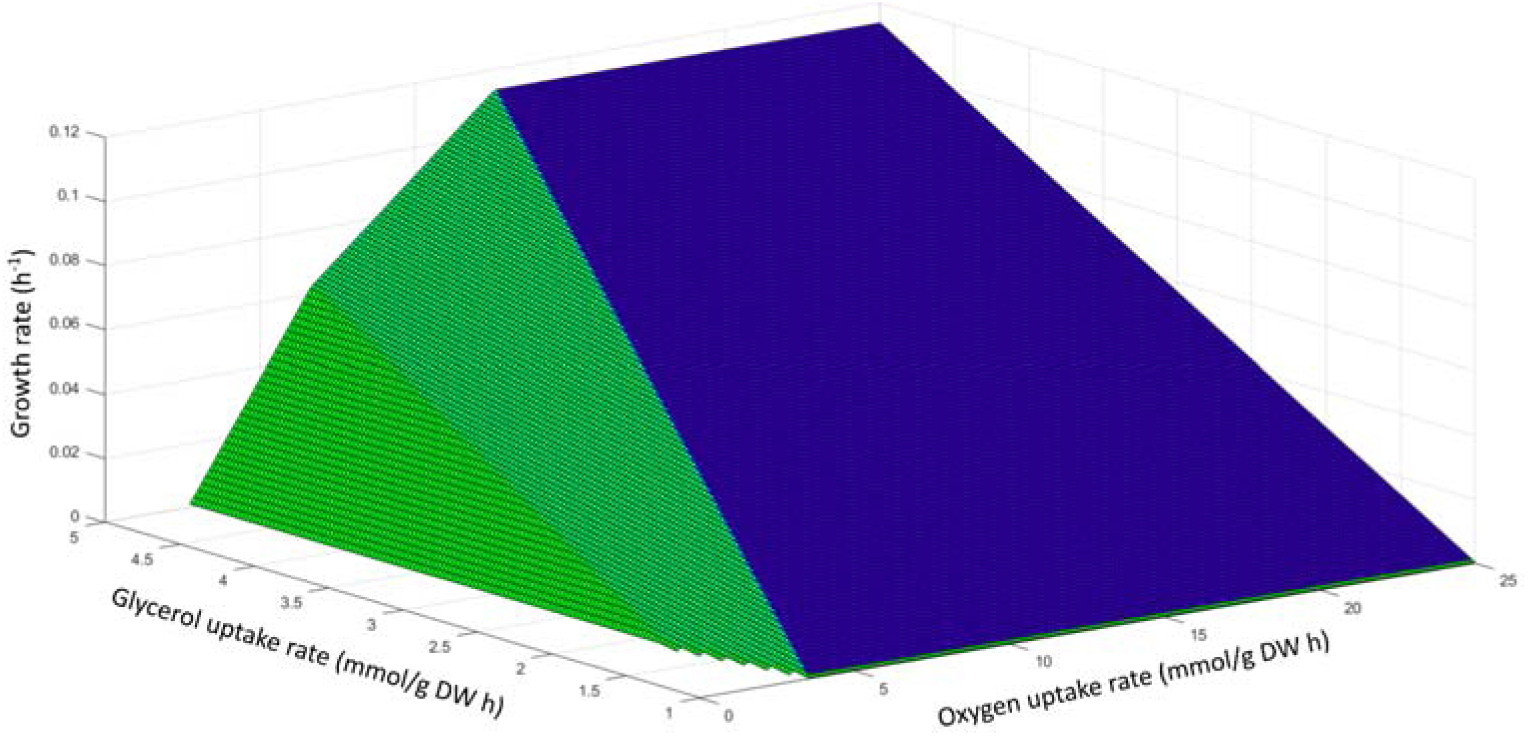
Effects of varying the glycerol and oxygen uptake rates on growth.

**Figure S4.**
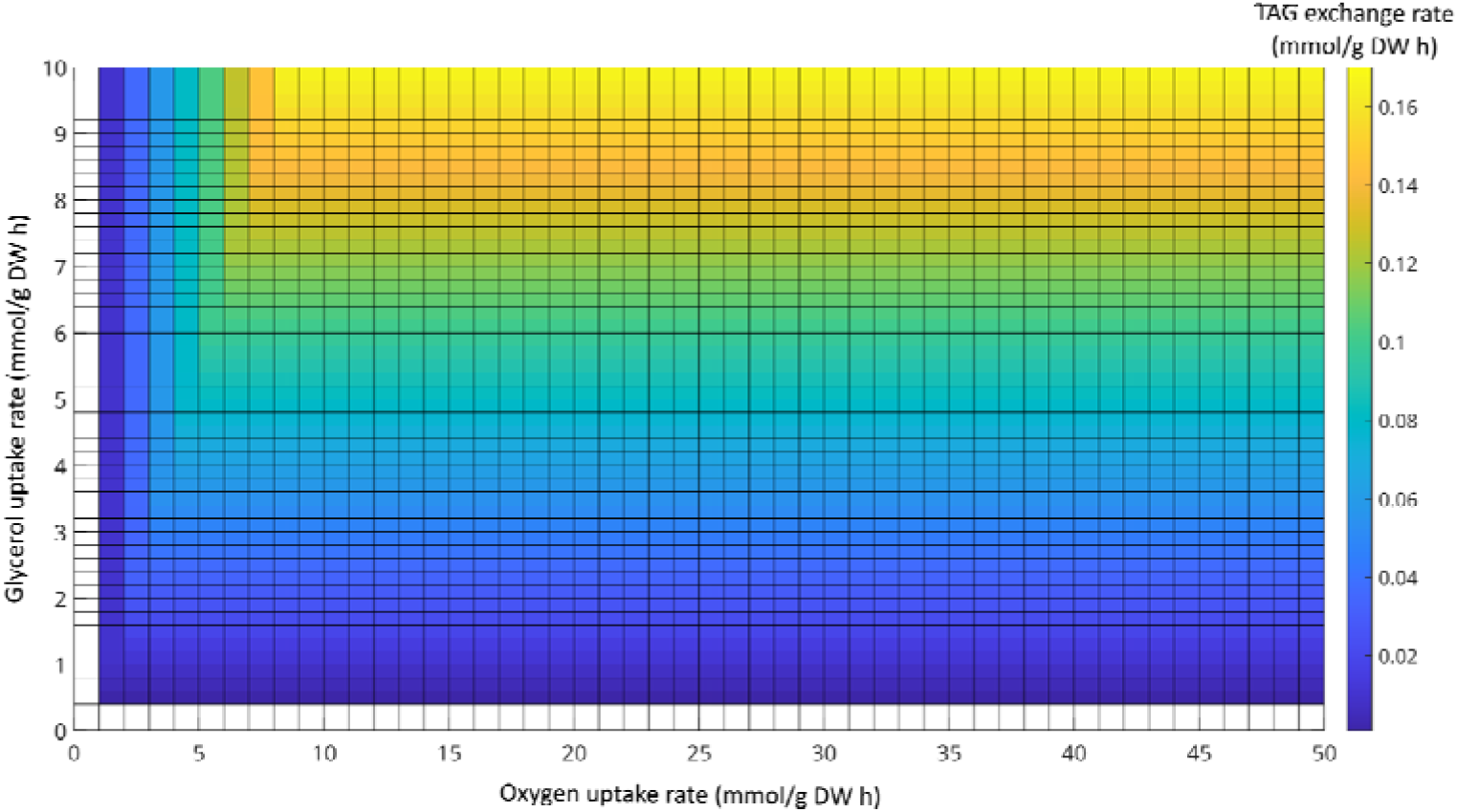
Effects of varying the glycerol and oxygen uptake rates on TAG exchange.

**Figure S5.**
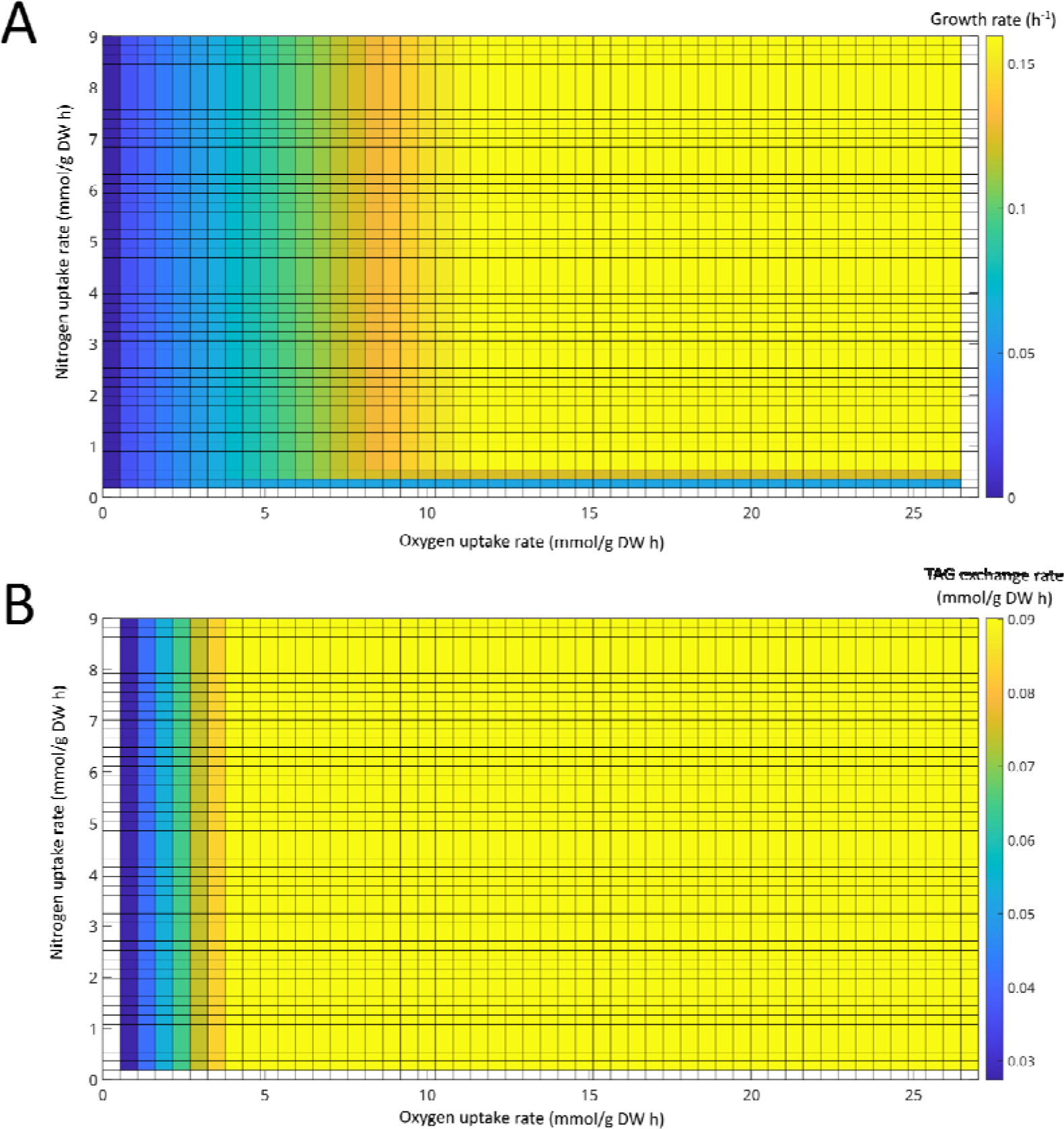
Effects of varying the nitrogen and oxygen uptake rates on (A) growth and (B) TAG exchange rates fixing glucose as the carbon source.

**Figure S6.**
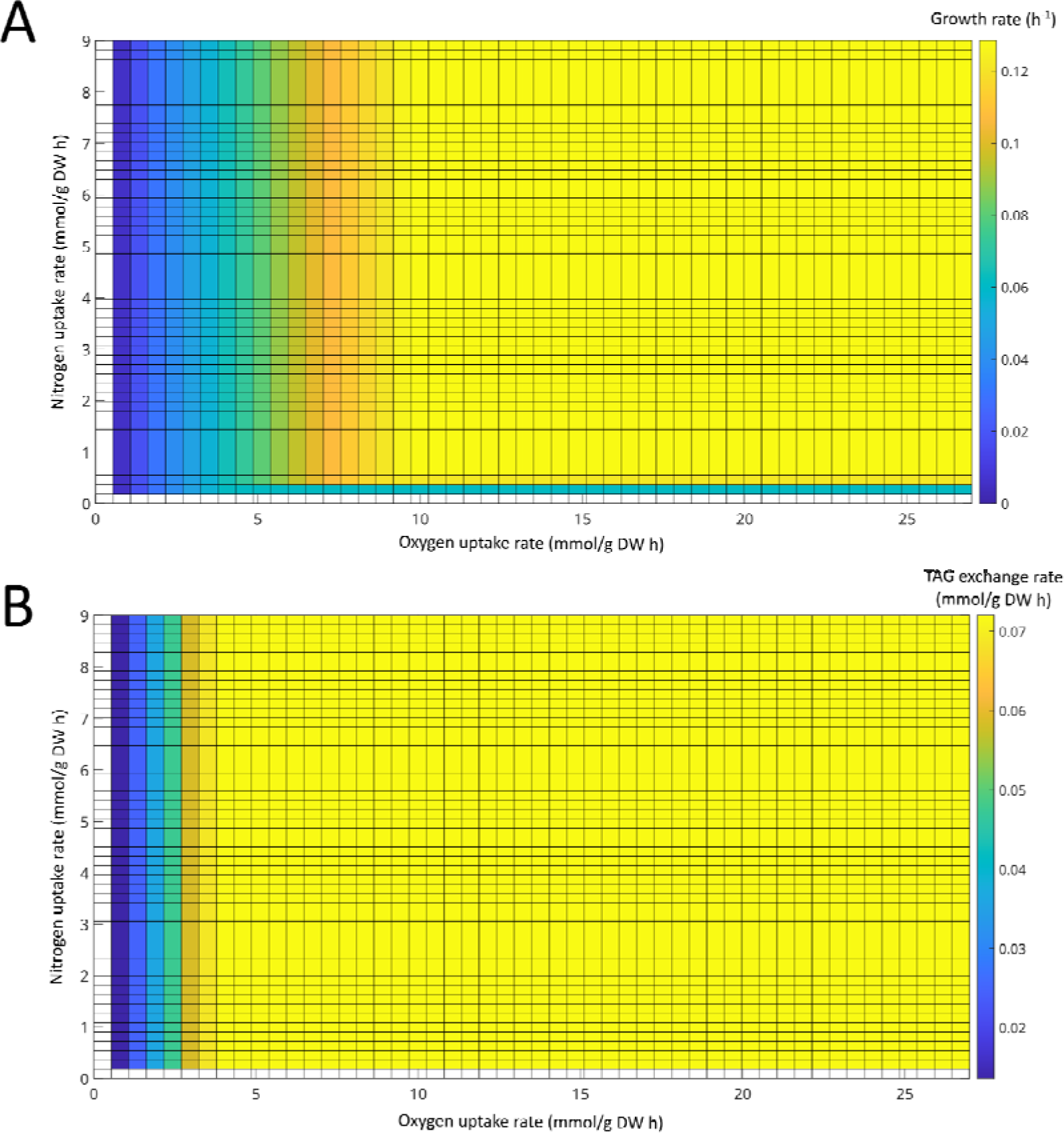
Effects of varying the nitrogen and oxygen uptake rates on (A) growth and (B) TAG exchange rates fixing xylose as the carbon source

**Figure S7.**
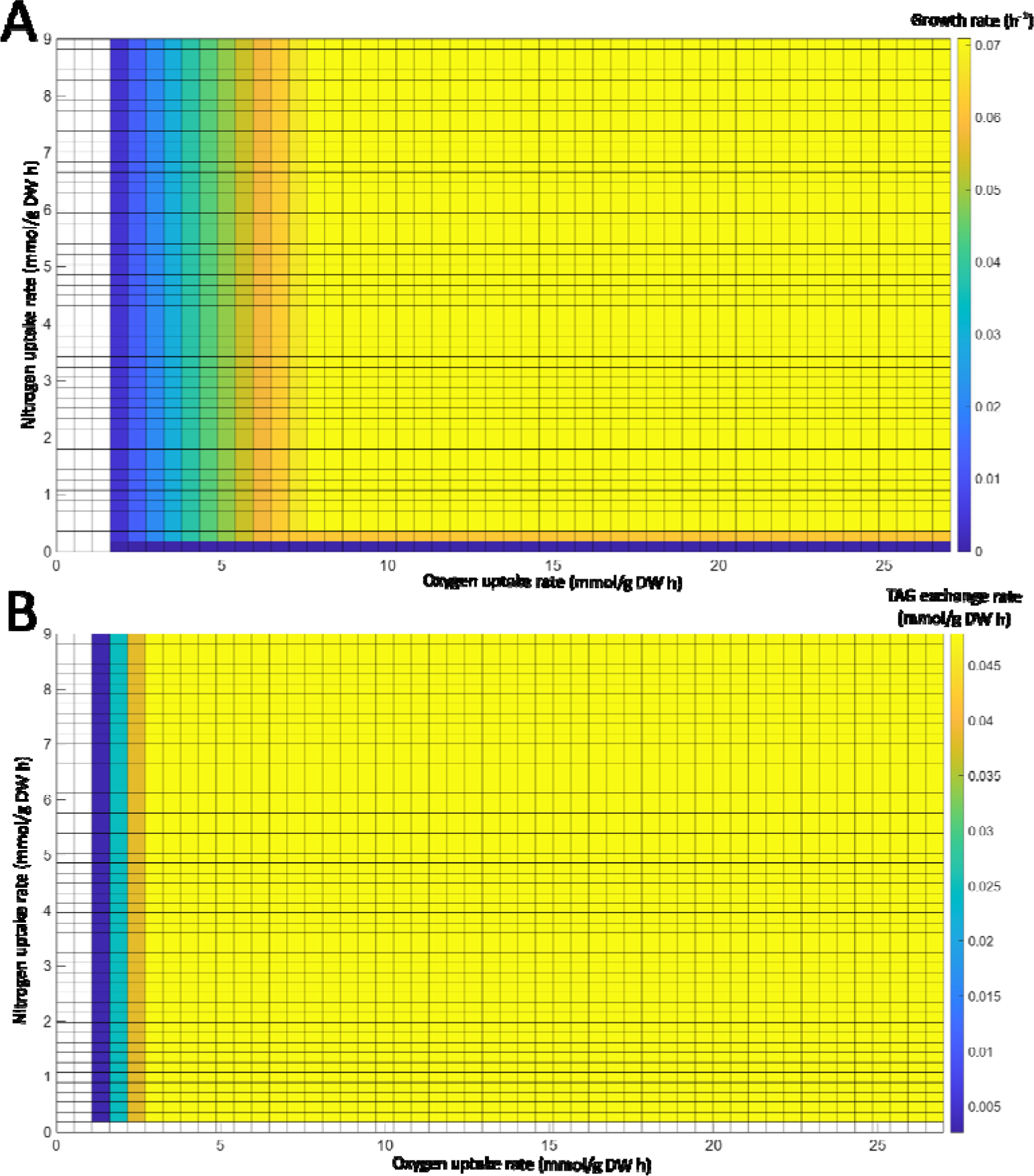
Effects of varying the nitrogen and oxygen uptake rates on (A) growth and (B) TAG exchange rates fixing glycerol as the carbon source.

## Notes

https://github.com/LabFisUFV/lista-GEM

https://doi.org/10.5281/zenodo.8367982

### Competing Interest Statement

The authors have declared no competing interest.

